# The viral oncoproteins Tax and HBZ reprogram the cellular mRNA splicing landscape

**DOI:** 10.1101/2021.01.18.427104

**Authors:** Charlotte Vandermeulen, Tina O’Grady, Bartimee Galvan, Majid Cherkaoui, Alice Desbuleux, Georges Coppin, Julien Olivet, Lamya Ben Ameur, Keisuke Kataoka, Seishi Ogawa, Marc Thiry, Franck Mortreux, Michael A. Calderwood, David E. Hill, Johan Van Weyenbergh, Benoit Charloteaux, Marc Vidal, Franck Dequiedt, Jean-Claude Twizere

## Abstract

While viral infections are known to hijack the transcription and translation of the host cell, the extent to which encoded viral proteins coordinate these perturbations remains unclear. Here we demonstrate that the oncoviral proteins Tax and HBZ interact with specific components of the spliceosome machinery, including the U2 auxiliary factor large subunit (U2AF2), and the complementary factor for APOBEC-1 (A1CF), respectively. Tax and HBZ perturb the splicing landscape in T-cells by altering cassette exons in opposing manners, with Tax inducing exon inclusion while HBZ induces exon exclusion. Among Tax- and HBZ-dependent splicing changes, we identify events that are also altered in Adult T cell leukemia (ATL) patients, and in well-known cancer census genes. Our interactome mapping approach, applicable to other viral oncogenes, has identified spliceosome perturbation as a novel mechanism coordinately used by Tax and HBZ to reprogram the transcriptome.

**Highlights:**

- Tax and HBZ interact with RNA-binding proteins as well as transcription factors
- HTLV-1 encoded proteins Tax and HBZ alter the splicing landscape in T-cells
- Tax and HBZ expression affect alternative splicing of 33 and 63 cancer genes, respectively
- Opposing roles for Tax and HBZ in deregulation of gene expression

**Graphical abstract:** 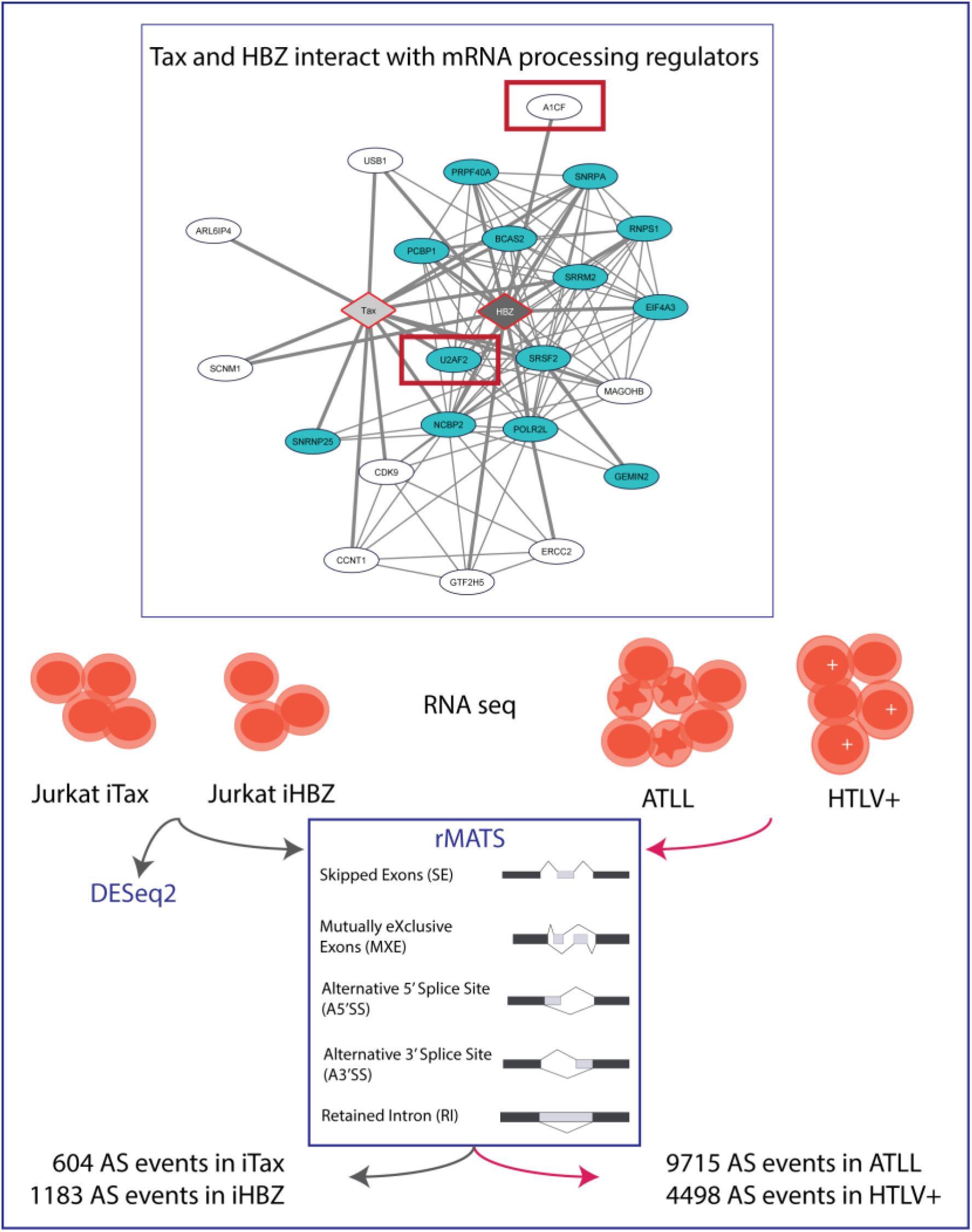

## INTRODUCTION

The ability of a retrovirus to transform its host cell was originally attributed to integration of retroviral DNA into the host cell’s genome. This integration allowed the discovery of cellular oncogenes and related cellular signaling pathways such as the SRC (Hunter and Sefton, 1980), EGFR (Downward et al., 1984), MYC (Hayward et al., 1981), RAS (Der et al., 1982), and PI3K pathways (Chang et al., 1997). However, no universal model has been developed to explain oncogenic transformation as a phenotypic result of retroviral integration. The only transforming retrovirus identified in humans to date is the human T-cell leukemia virus type 1 (HTLV-1), which causes adult T-cell leukemia/lymphoma (ATLL). ATLL has a long latency period of approximately 20-60 years, which suggests the occurrence of rare and complex genomic changes during disease progression (Kataoka et al., 2018, 2015). Proviral integration sites for HTLV-1 are enriched in cancer driver genes, resulting in altered transcription of those genes (Rosewick et al., 2017). Additional genomic changes have also been observed in distant genes that play a role in various mechanisms of T-cell signaling (Kataoka et al., 2018, 2015). However, the key initial drivers of ATLL are the viral proteins Tax and HBZ, which can independently induce leukemia in transgenic mouse models (Hasegawa et al., 2006; Satou et al., 2011).

Studies of Tax and HBZ have been summarized in several reviews (Baratella et al., 2017; Boxus et al., 2008; Yasunaga and Matsuoka, 2018). These data have been compiled into a KEGG pathway (hsa05166) which highlights the ability of Tax and HBZ to interfere with at least one component of each of the twelve signaling pathways regulating the three cancer core processes: cell fate, cell survival, and genome maintenance (Vogelstein et al., 2013). The effects of Tax and HBZ are mediated primarily via protein-protein interactions, and positive or negative transcriptional regulation (Boxus et al., 2008; Simonis et al., 2012). Tax and HBZ often act in opposing directions in order to control the host’s immune response and sustain long-term malignant transformation (Sugata et al., 2015; Yasuma et al., 2016).

Our molecular understanding of genomics and transcriptomics deregulation following HTLV-1 infection has come from studying ATLL patient samples (Kataoka et al., 2015; Thénoz et al., 2014). Because Tax and HBZ show different expression kinetics during ATLL progression, it has remained challenging to systematically analyze the relative contribution of each viral protein in reprogramming the host cell’s transcriptome and proteome. Here, we carried out a systematic identification of the interactome networks between Tax/HBZ and cellular regulators of gene expression. We then measured the effects of Tax and HBZ on gene expression at both the transcriptional and post-transcriptional levels. Integration of these interactome and transcriptome datasets provides mechanistic insights into HTLV-1 infection-associated alternative splicing events.

## RESULTS

### A comparative interactome of Tax and HBZ with cellular host proteins

Previous studies have shown that Tax and HBZ viral proteins control viral gene expression by competing for binding to key transcriptional factors of the CREB/ATF pathway, and coactivators CBP and p300 (Figures 1A-B), complexes that are also specifically targeted by other viral oncoproteins including high-risk human papillomavirus (HPV) E6 proteins (Rozenblatt-Rosen et al., 2012). In the present study, we aimed at providing an unbiased map of protein-protein interactions (PPIs) established by Tax and HBZ viral oncoproteins with cellular gene expression regulators, including transcription factors (TFs) and RNA-binding proteins (RBPs) (Figure 1C). Our rationale is that exploring more broadly the similarities and differences of Tax and HBZ interactomes with TFs and RBPs would provide global and specific insights on how viral oncogenes cooperate in the initiation and maintenance of cancer.

**Figure1.**
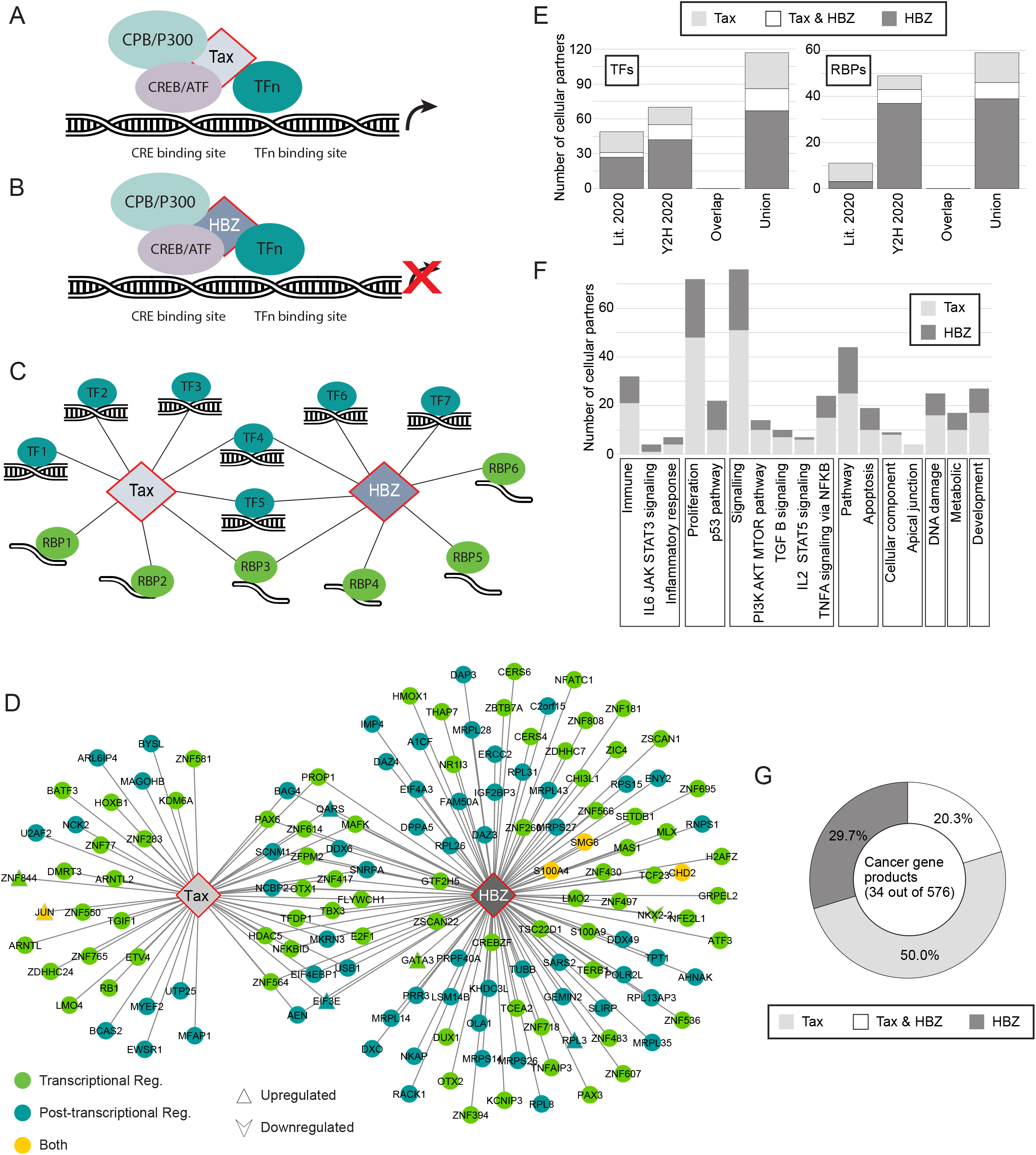
A comparative interactome network of Tax and HBZ with cellular host genes. **(A)** Schematic representation of positive interaction between Tax, CREB/ATF and CBP/P300 on the viral promoter. **(B)** As in (A) but negative interaction driven by HBZ. **(C)** Illustrative network diagram showing interactions between Tax and HBZ with cellular transcription factors (TF) or RNA-binding proteins (RBP). **(D)** Y2H Interactome map of Tax and HBZ with host TRs (blue), PTRs (green) or both TR/PTR (yellow). **(E)** Graph showing the number of RBPs and TFs interacting with Tax and HBZ in different datasets. **(F)** As in (E) but number of coding genes that are parts of a GSEA hallmark category. **(G)** Percentages of Cancer gene products interacting with Tax and HBZ. See also Figures S1, S2 and Tables S1 and S3.

Based on Gene Ontology and literature curation, we first established a library that covers 3652 ORFs encoding 2089 transcriptional and 1827 post-transcriptional regulators, including known DNA- and RNA-binding proteins (Figure S1A). Secondly, we assembled a mini-library of different *Tax* and *HBZ* clones with previously described functional domains (Figure S1B, Table S1; Boxus et al., 2012, 2008; Clerc et al., 2008; Forlani et al., 2016; Giam and Semmes, 2016; Hivin et al., 2005; Pise-Masison et al., 2000; Takachi et al., 2015). We then tested binary interacting pairs between viral products and human TFs and RBPs using our well-established binary interactome mapping strategy employing primary screening by yeast two-hybrid (Y2H) assays, retesting by Y2H and validation using an orthogonal protein complementation assay (Luck et al., 2020; Roland et al., 2014; Bergiers et al., 2014; Dreze et al., 2010). This interactome search space encompassing ∼95,000 binary combinations, allowed identification of 53 and 116 Tax and HBZ cellular partners, respectively (Figures 1D-E and S1, Table S1). Interestingly, we observed a highly significant overlap between gene expression regulators, either TFs or RBPs, interacting with Tax and HBZ (Figure 1D, 25 shared interactors, Fisher test: p<2.2e^-16^). We next used 90 interacting pairs in a validation experiment using an orthogonal assay, the *Gaussia princeps* protein complementation assay (GPCA) (Cassonnet et al., 2011). The validation rate was 83%, demonstrating the high quality of our dataset (Y2H_2020) that represents an increase of 32% and 51.6% of known TFs and RBPs interacting with Tax, respectively (Figures 1E and S1D-E). Compared to Tax, fewer interactions were available in the literature for HBZ, and our result represents a substantial increase of 65.5% and 93.6 % of TFs and RBPs interacting with HBZ, respectively (Figure 1E).

We expanded our catalog of systematically determined binary Tax and HBZ interactions with host proteins (Y2H_2020) with high quality interactions reported in the literature (Lit_2020) (Figures 1E and S2). The union of Tax-host and HBZ-host interactomes contains 258 and 160 interacting partners, respectively (Figure S2 and Table S1). Of interest, TFs represent 38% and 59%, while RBPs count for 16% and 38% of Tax and HBZ host interactors, respectively (Figure S2 and S1).

To classify Tax and HBZ interacting partners in functional categories we examined their repartition in the MSigDB hallmark gene sets (Liberzon et al., 2015) (Figure 1F). We focused on five categories of specific gene sets (Immune, Proliferation, Signaling, Pathway and Cellular component) and obtained a constructive view of the functional distribution of Tax and HBZ partners. The most significantly enriched gene set signatures were “Proliferation” and “Signaling pathways”, for both Tax and HBZ (Figure 1F). While HBZ appears to specifically target the IL-6-JAK-STAT3 pathway, cancer gene products of the PI3K-AKT-mTOR, TGF-β and IL-2-STAT5 pathways are more enriched in the Tax interactome (Figure 1F). Further highlighting the impact of Tax and HBZ in the initiation and maintenance of cancer, these viral proteins exhibit significantly more PPIs (at least 20 times) than the average “degree” (number of interactors) of known cancer gene products (Rolland et al. 2014) or other tumor virus proteins (Rozenblatt-Rosen et al., 2012).

Altogether these results provide strong evidence that Tax and HBZ perturb the cell host through similar and differential associations with transcriptional and post-transcriptional regulators.

### A comparative analysis of transcriptomic changes associated with Tax and HBZ expression

Based on the observation that Tax and HBZ target numerous host DNA- and RNA-binding proteins, it is anticipated that a significant number of host gene expression changes, indispensable for ATLL pathogenesis, are driven by these interactions. However, since Tax and HBZ are differentially expressed *in vivo*, and compete for binding to key transcriptional regulators, it is impractical to systematically compare and identify endogenously specific Tax- and HBZ-induced changes in infected cells or a single cell line exogenously expressing both Tax and HBZ. Thus, we generated a homogeneous inducible cellular system, which consists of two Jurkat T cell lines, Jurkat-iTax and Jurkat-iHBZ, expressing either Tax or HBZ from a doxycycline-inducible promoter (Figure 2A). We then performed high-throughput RNA sequencing (RNA-seq) of the two cell lines and analyzed differentially expressed genes (DEGs), and alternative splicing events (ASEs) associated with Tax or HBZ expression (Figure 2A). We confirmed the induction of Tax and HBZ (Figures 2B and S3A), which are associated with increased expression of their common up- or down-regulated target genes *GATA3* (Figure 2B) or *SYT4* (Figure S3C), respectively (Blumenthal et al., 1999; Kataoka et al., 2015; Sugata et al., 2016), and differentially regulated *STAT5A* (Figure S3B) (Nakamura et al., 1999; Zhao et al., 2011).

**Figure 2.**
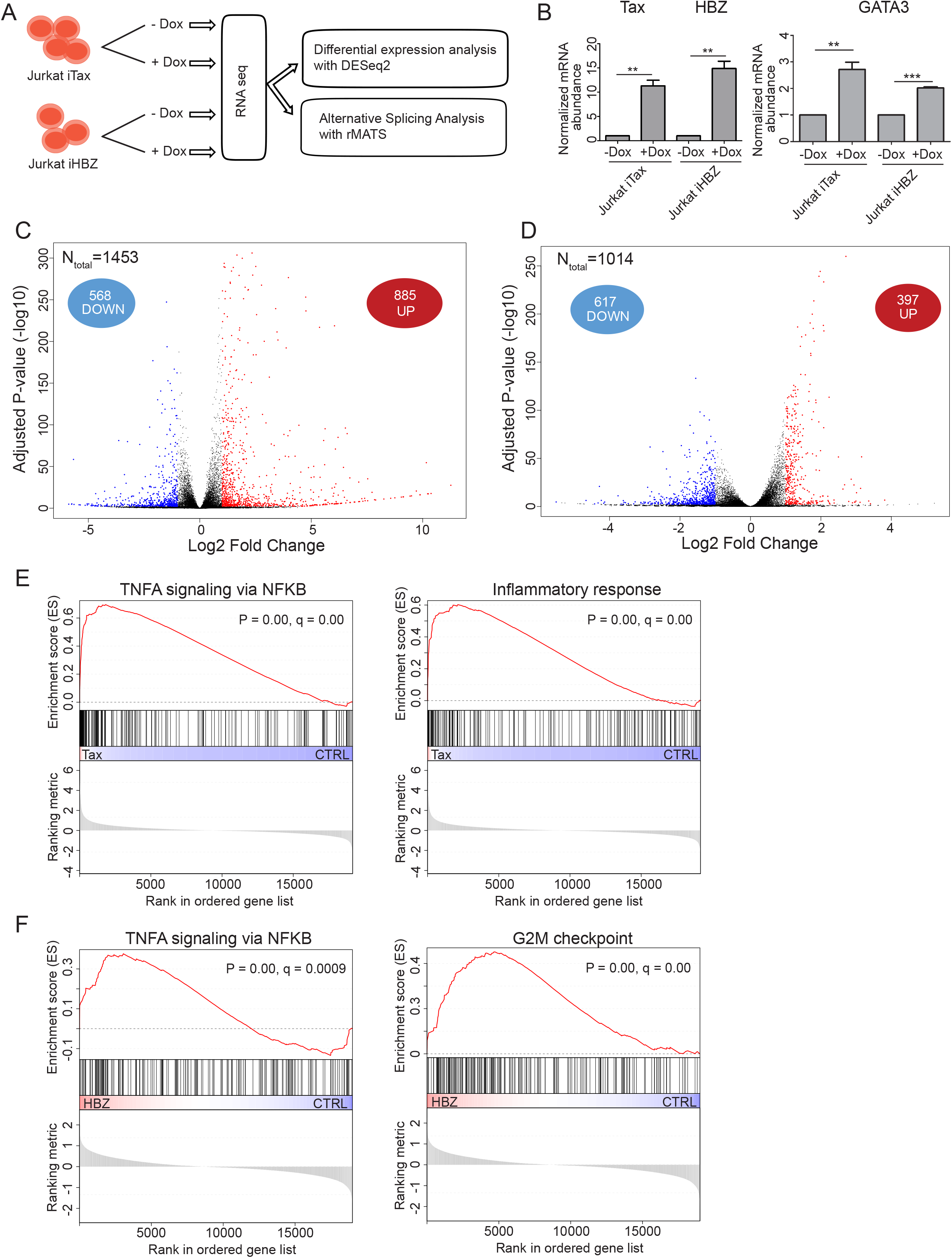
A comparative analysis of transcriptomic changes associated with Tax and HBZ expression. **(A)** Overview of the experimental design for analysis of abundance and splicing alterations upon Tax and HBZ expression in Jurkat cells. **(B)** Tax and HBZ expression was analyzed by qRT-PCR in Jurkat cells after induction with doxycycline (left). Overexpression of GATA3 in Jurkat-iTax and Jurkat-iHBZ cell lines (right). **(C-D)** Volcano plot of significantly up-regulated genes (red dots) and down-regulated genes (blue dots) upon induction of Tax (C) or Flag-HBZ (D) in Jurkat cells. **(E-F)** Examples of significantly enriched gene sets by GSEA in Jurkat-iTax (E) cells and Jurkat-iHBZ (F) cells. See also Table S2.

Comparative analysis with control cells revealed 1453 and 1014 genes that were differentially expressed upon inducing Tax or HBZ expression, respectively (FDR adjusted p< 0.01 and |log2 fold-change|≥1, Figures 2C and 2D and Tables S2). Consistent with its well-known function as a transcriptional activator (Franklin et al., 1993; Hirai et al., 1992), we found that Tax expression caused up-regulation of 885 genes and down-regulation of 568 genes (Figure 2C). In contrast, expression of HBZ was associated with up-regulation of 397 genes and down-regulation of 617 genes (Figure 2D). Several studies found that HBZ can act as a transcription repressor (Basbous et al., 2003; Clerc et al., 2008; Lemasson et al., 2007; Takiuchi et al., 2017; Tanaka-Nakanishi et al., 2014; Wurm et al., 2012), in agreement with our finding that 61% of the differentially regulated genes in Jurkat-iHBZ were down-regulated. This finding is also supported by our confocal and transmission electron microscopy observations showing that HBZ-expressing cells have larger nuclear speckles (Figures S3D and S3E), which is often associated with decreased Pol II−mediated transcription (Didichenko and Thelen, 2001; Lamond and Spector, 2003).

Gene Set Enrichment Analysis (Subramanian et al., 2005) identified TNF-α signaling via NF-kB, as a specific pathway enriched both in Tax- and HBZ-expressing cells (Figures 2E and 2F). In contrast, inflammatory response was specifically up-regulated in Tax-expressing cells whereas cell cycle G2M checkpoint genes were specifically up-regulated in HBZ expressing cells (Figures 2E and 2F and Table S2). Interestingly, the extent of co-regulated genes by both viral proteins was highly significant (28% for Tax and 41% for HBZ, empirical P < 0.0001). These include 405 genes whose expression was altered in the same direction (149 up-regulated and 256 down-regulated) and only 12 genes whose expression was altered in opposite directions (Table S2). Despite their differential expression *in vivo*, our transcriptomic changes analysis confirms the notion highlighted above, that Tax and HBZ viral proteins drive a number of overlapping molecular perturbations.

### Impact of Tax and HBZ expression on cellular gene alternative splicing events

Analyses of Tax and HBZ host cell perturbations have predominately focused on transcriptional defects. However, from our interactome data, the interacting proteins annotated as “post-transcriptional regulators” represent 54% and 44% of Tax and HBZ partners involved in gene expression regulation, respectively (Figure S2). To assess the impact of Tax and HBZ on the cellular splicing landscape, we statistically computed splicing events using the rMATS software (Figure 3A, Shen et al., 2014). We focused on 5 types of alternative splicing: Skipped Exons (SE) or cassette exons, Mutually Exclusive Exons (MXE), alternative 3’ Splice Sites (3’SS), alternative 5’ Splice Sites (5’SS) and Retained Introns (RI) (Shen et al., 2014), which we quantified as Percent Spliced In (PSI), (Figure 3A and Table S3). Our analysis identified 604 Tax- and 1183 HBZ-regulated alternative splicing events (ASEs) corresponding to 447 and 839 genes, respectively (Figure 3B). The majority of Tax and HBZ-dependent events were cassette exons (65.5% and 75.9% for Tax and HBZ, respectively) (Figure 3B).

**Figure 3.**
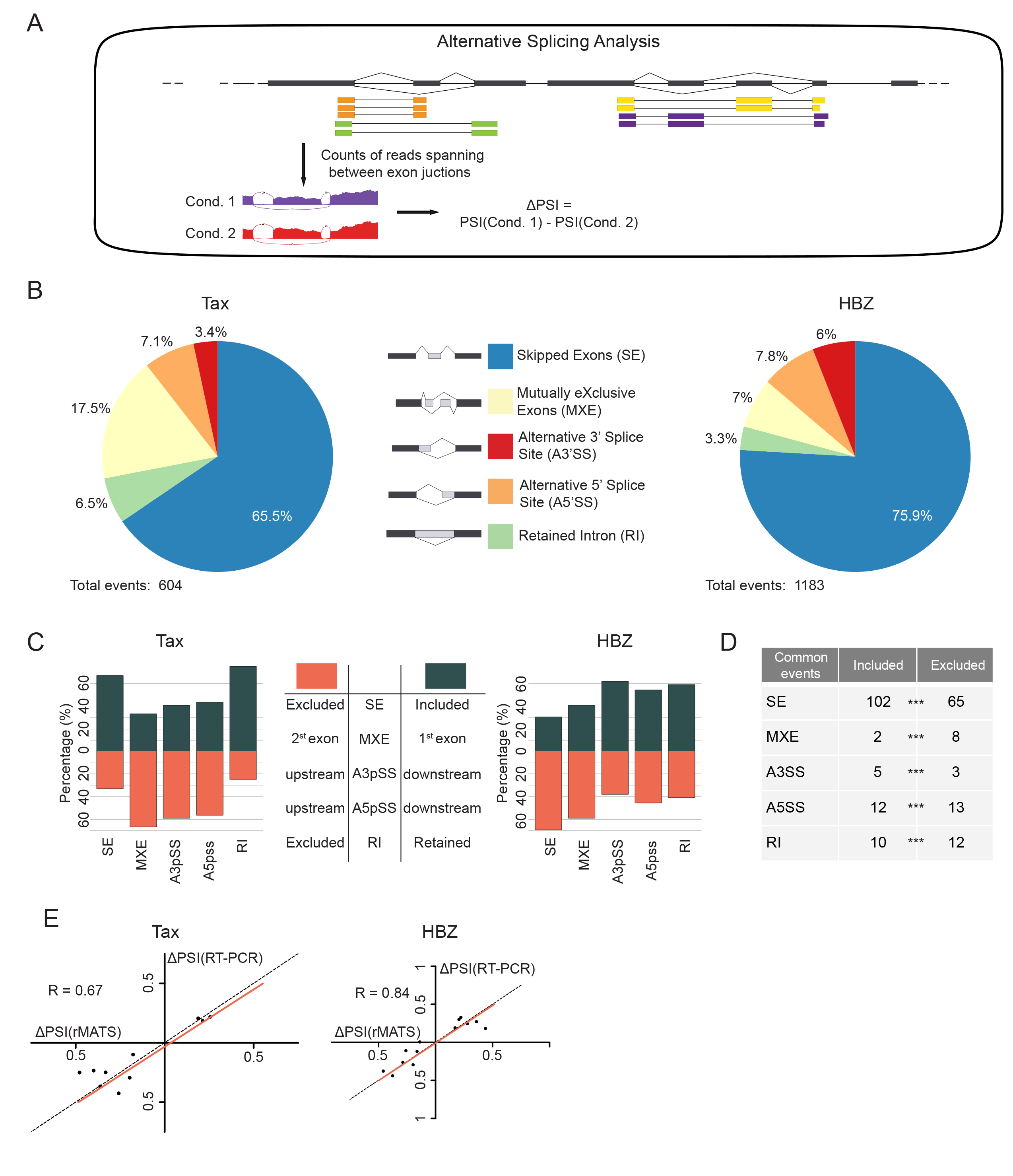
Impact of Tax and HBZ expression on cellular gene alternative splicing events. **(A)** rMATS analysis to determine alternative splicing between conditions. **(B)** Splicing profiles of ASEs detected in Jurkat cells expressing Tax (left) or Flag-HBZ (right). SE = Skipped Exon, MXE = Mutually Exclusive Exons, A3’SS = Alternative 3’ Splice Site, A5’SS = Alternative 5’ Splice Site, RI = Retained Intron. **(C)** Exclusion or inclusion of ASEs observed in Tax and Flag-HBZ expressing conditions. **(D)** Number of shared ASEs between Jurkat cells expressing Tax or Flag-HBZ. **(E)** Graph showing the Spearman correlation between the ΔPSI values obtained with RNA-seq and RT-PCR. See also Figure S3 and Table S3.

We validated the RNA-seq analysis by qRT-PCR for 10 Tax-dependent and 13 HBZ-dependent SEs (10/10 for Tax, 12/13 for HBZ, Figures 3E). Interestingly, Tax and HBZ had globally opposite effects on the cellular splicing landscape (Figures 3C). This was particularly true for SE where most regulated exons (66.9%) showed increased inclusion upon Tax expression. In contrast, the majority of HBZ-regulated cassette exons (69.3%) were more likely to be skipped. However, we also found a significant overlap between the different Tax- and HBZ-regulated ASEs: 167 SE (P = 3.51e-130), 8 A3’SS (P = 4.17e-12), 25 A5’SS (P =3.24e-31), 10 MXE (P = 2.10e-09) and 22 RI (P = 7.15e-20) (Figure 3D).

As observed for differentially expressed genes, a number of SE exons were similarly regulated by Tax and HBZ expression (102 inclusion and 65 exclusion events in both Tax and HBZ expressing cells, Figure 3D). Taken together our analysis shows that although Tax and HBZ have globally opposite effects on the host splicing landscape, they share some common target exons that are similarly affected.

### Splicing targets of Tax and HBZ are enriched for cancer related genes

Gene Ontology (GO) enrichment analysis of splicing targets revealed significant enrichment of several GO terms in HBZ-regulated genes, whereas no significant enrichment was detected for Tax splicing targets. For HBZ, enriched categories were mostly related to RNA regulation, especially RNA splicing (Table S3. We further investigated the functions of specific spliced genes in cells expressing Tax or HBZ by examining their repartition in the MSigDB hallmark gene sets built by GSEA (Liberzon et al., 2015) (Figure 4A). A significant overlap was also found between genes corresponding to Tax or HBZ splicing pre-mRNA targets and known cancer census genes (ncg.kcl.ac.uk and cancer.sanger.ac.uk) (P =0.02247 for Tax and P = 0.001742 for HBZ). Pre-mRNA of 33 and 63 cancer genes were identified as splicing targets of Tax and HBZ, respectively (Table S3). Among these, 10 were similarly deregulated by Tax and HBZ (*CHCHD7, EIF4A2, NF2, POLG, PTPRC, UBR5, ABI1, BCLAF1, FLNA* and *NSD1*). These include *PTPRC*, coding for CD45, a transmembrane protein tyrosine phosphatase, which is known to be alternatively spliced upon T-cell differentiation and induces a switch from naive (CD45RA) to memory T-cells (Hermiston et al., 2003; Heyd et al., 2006; Lynch and Weiss, 2000; Mazurov et al., 2012).

**Figure 4.**
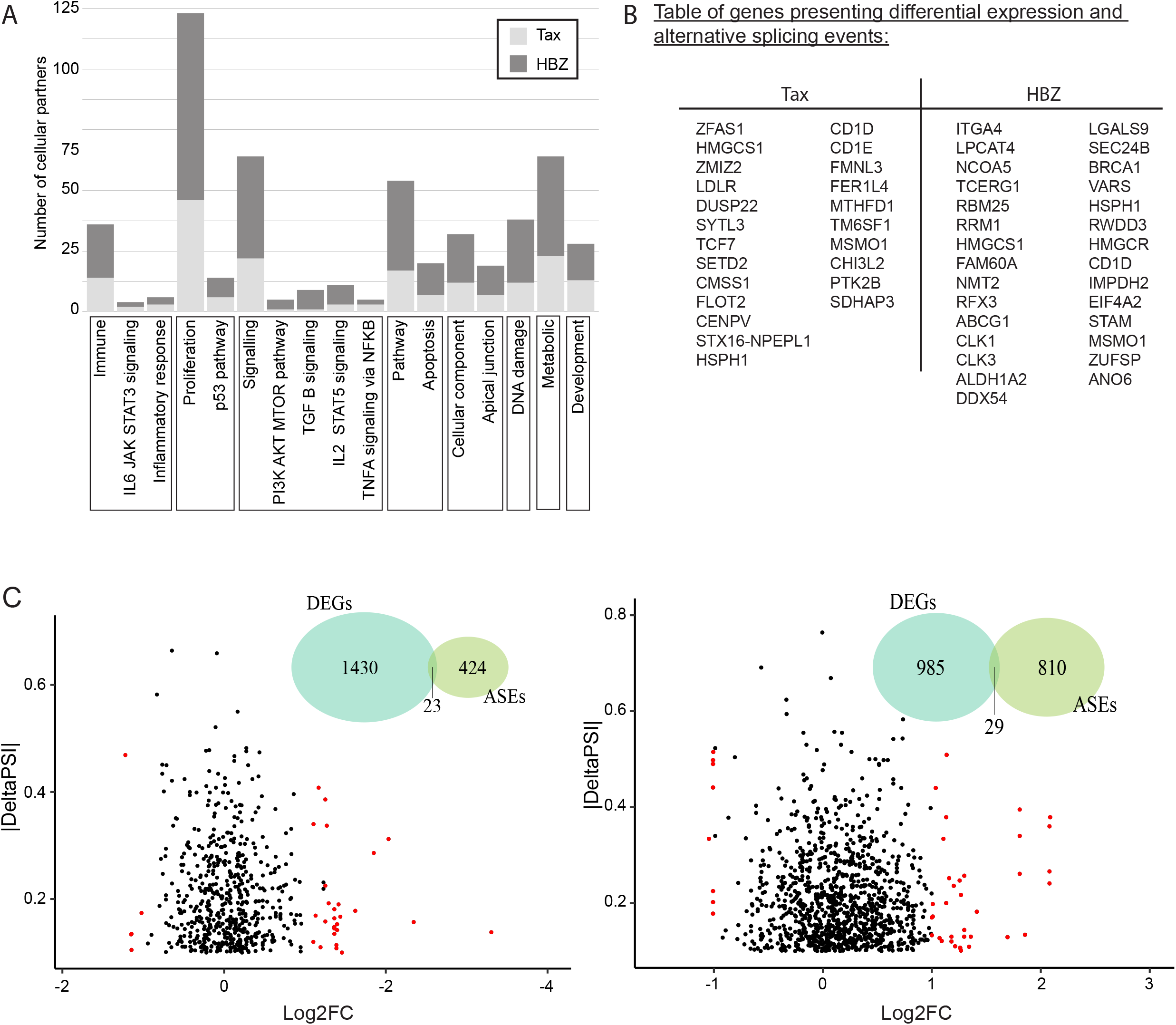
Splicing targets of Tax and HBZ are enriched for cancer related genes. **(A)** Alternatively, spliced (AS) genes in iTax or iHBZ cells, which are part of a GSEA hallmark category. Cancer genes are in red. **(B)** Table of genes with DE and AS in Tax or HBZ expressing cells **(C)** ΔPSI of AS genes and their differential gene expression (DE) upon Tax (left) or Flag-HBZ (right) expression. AS and DE genes are shown in red. Venn diagrams show up- or down-regulation upon Tax or HBZ expression (blue) and genes with ASEs (green).. See also Figure S4 and Table S3.

Another interesting observation here is the fact that only a small number of genes (23 and 29 for Tax or HBZ, respectively) presenting ASEs were differentially expressed (Table S3), suggesting that different sets of genes are regulated at the transcriptional and Mrna splicing levels (Figures 4B and 4C).

### Splicing events in ATLL patients explained by Tax or HBZ expression

To determine whether ASEs detected in Tax or HBZ expressing cells could be relevant for HTLV-1 infection and leukemogenesis, we interrogated RNA-seq expression data obtained from 35 ATLL samples, 3 samples from HTLV-1 carriers without ATLL and 3 samples from healthy volunteers (Kataoka et al., 2015). We detected 4497 ASEs (in 2343 genes) between HTLV-1 carriers and healthy volunteers, while 9715 events (in 2737 genes) were differentially affected in the ATLL patients compared to healthy volunteers. Among these, cassette exons and mutually exclusive exons accounted for a large majority of ASEs (Figures 5A and 5B), as observed for cells expressing Tax or HBZ proteins (Figure 3B). Interestingly, we observed a significant overlap between ASEs detected in HTLV-1 carriers and ATLL patients (1637 ASEs on 1238 genes, SE P ∼0, A3’SS P = 8.026e-112, A5’SS P = 4.14e-118, MXE P = 1.40e-181, RI P = 5.01e-19), suggesting that some ASEs may initiate disease progression and persist during ATLL. Similar to our observations in Jurkat cells expressing Tax, alternative splicing of cassette exons (SE events) was skewed towards inclusion for both HTLV-1 carriers and patients with ATLL (Figure 5B).

**Figure 5.**
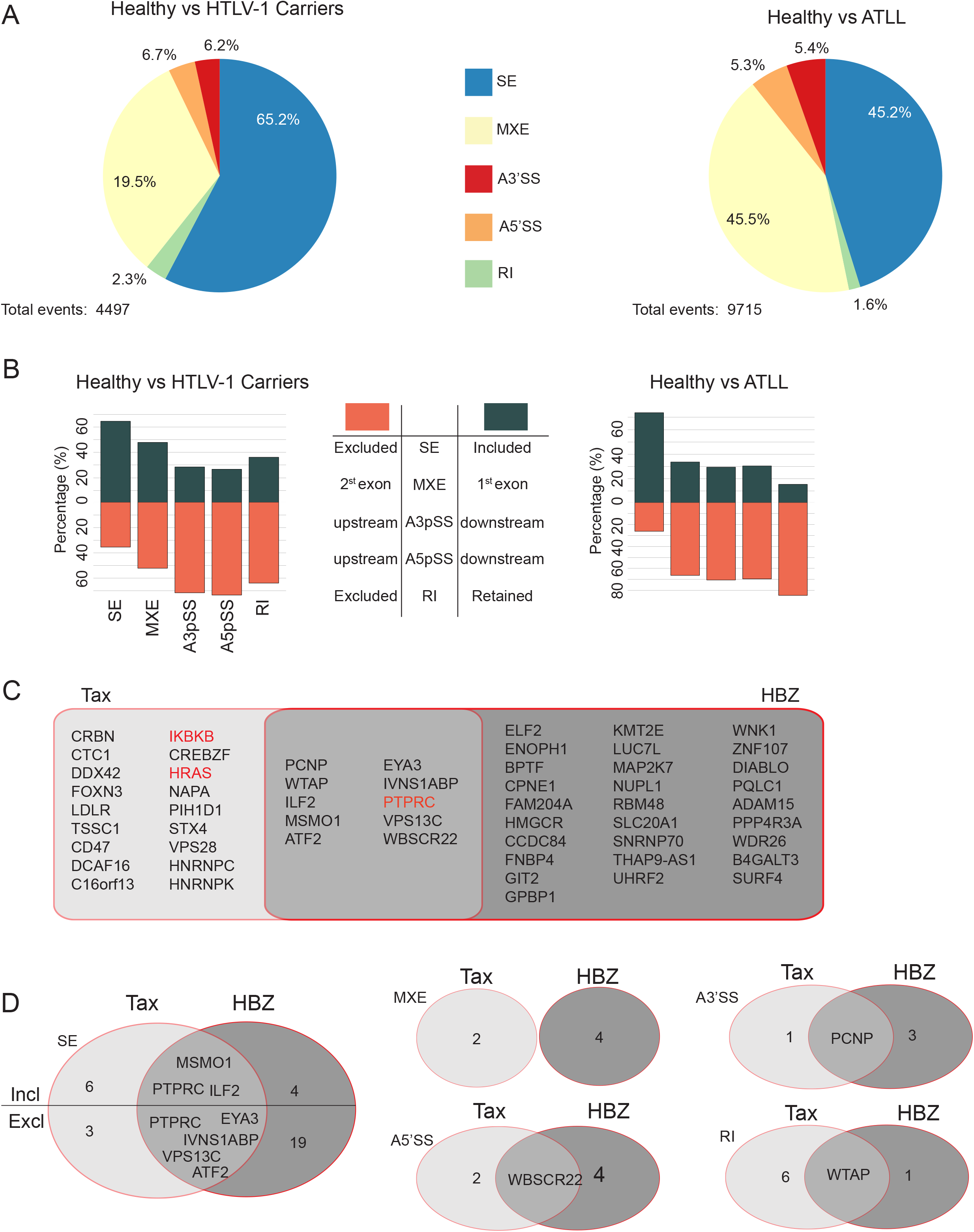
Splicing events in ATLL patients explained by Tax or HBZ expression. (A) ASEs detected in HTLV-1 carriers or patients with ATLL compared to healthy human peripheral CD4+ T cells. SE = Skipped Exon, MXE = Mutually Exclusive Exons, A3’SS = Alternative 3’ Splice Site, A5’SS = Alternative 5’ Splice Site, RI = Retained Intron (B) Exclusion or inclusion of ASEs observed in HTLV-1 carriers or ATLL. (C) Genes sharing similar ASEs in ATLL and Jurkat cells expressing Tax or Flag-HBZ. (D) Number of shared events. See also Table S3.

The most significant GO-term enrichment for genes presenting ASEs in ATLL patients was RNA binding (GO:0003723). However, other categories were also significantly enriched, such as cadherin binding, viral process and immune system process (Table S4). Thirty-one and 42 ASEs observed in cells expressing Tax or HBZ, respectively, were also present in patients with ATLL. Those events occurred on pre-mRNA of 56 genes including well-known cancer-related genes *PTPRC, IKBKB* and *HRAS*, and genes coding for transcription factors ILF2, ATF2and EYA3 (Figures 5 C-D). Notably, several ASEs on the *PTPRC* pre-mRNA coding for CD45 that were detected in Tax- and HBZ-expressing cells, were also present in patients with ATLL. These ASE changes occurred on *PTPRC* exons 4, 5, 6 and 7. More specifically, we observed a tendency for exons 5, 6 and 7 to be excluded, while exon 4 was more included (Figure 6A-D), suggesting that, compared to classical activated and memory T-cells (Lynch, 2004), ATLL cells exhibit a specific alternative splicing pattern on *PTPRC* pre-mRNA. Those exons are important for the production of diverse CD45 isoforms and therefore, the inclusion of exon 4 may represent a major mechanism used by Tax and HBZ to control T cell activation. Other examples include *ATF2* and *EYA3*, which present premature stop codons on spliced exons; and *MSM01*, which has a Nuclear Localization Signal (NLS) affected by splicing events (Figure S4). Altogether, our results demonstrate a global impact of Tax and HBZ expression on alternative splicing. Strikingly, Tax and HBZ targets show different splicing patterns in HTLV-1-infected individuals and ATLL patients, further emphasizing their global opposing effects on the deregulation of host genes.

**Figure 6.**
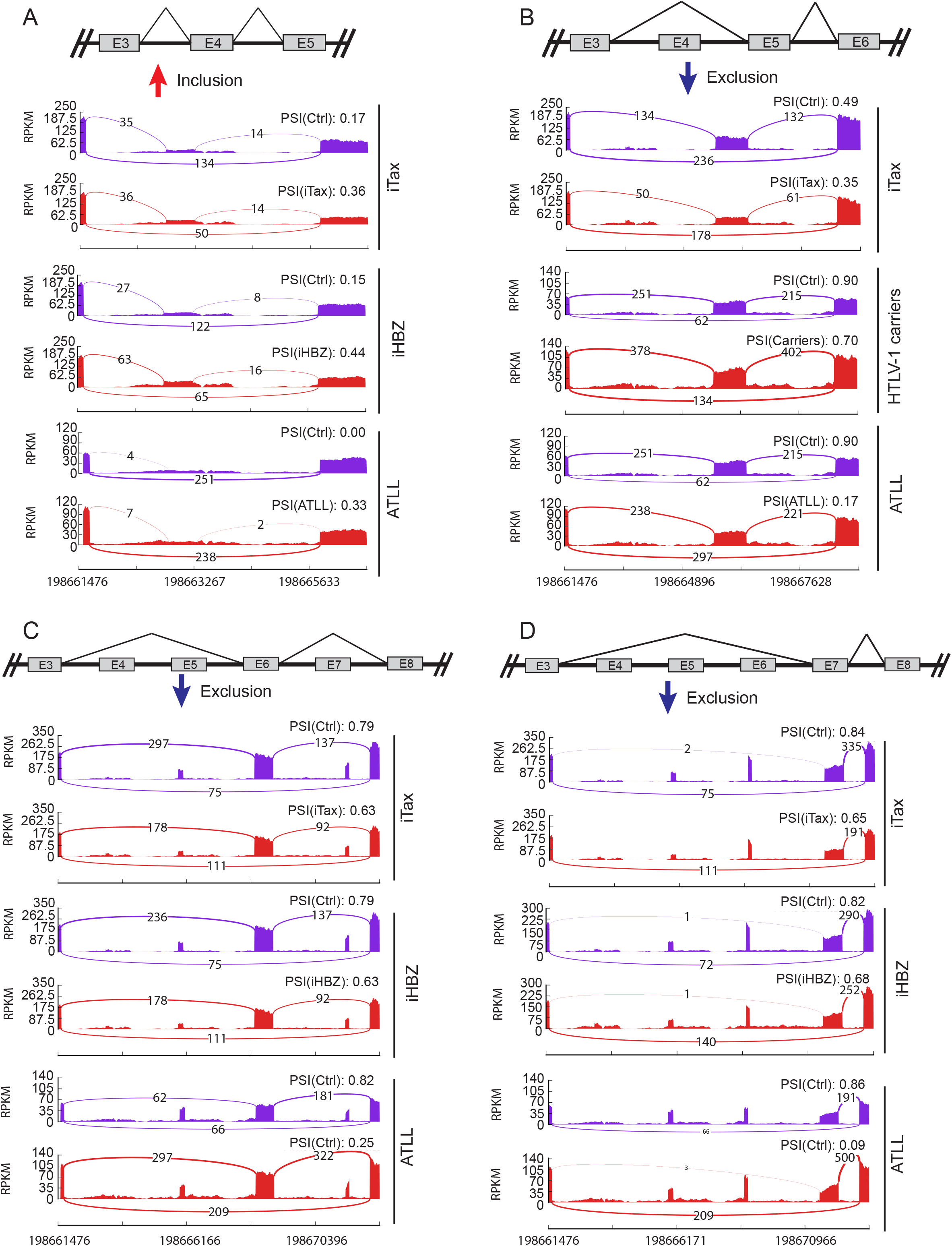
Alternative splicing events detected on exons 4 to 7 of PTPRC. Sashimi plots of significant ASEs detected on exons 4 to 7 of *PTPRC* gene in all analyzed conditions (indicated on the right). Splicing events are depicted above each sashimi plot, coordinates of regulated and flanking exons are indicated at the bottom. (A) Increased inclusion of exon 4. Decreased inclusion of exon 5 (B), 6 (C) and 7 (D). For ATLL, sashimi plot is a representative case. See also Table S3.

### Identification of RNA splicing-specific roles for Tax and HBZ proteins

Tax and HBZ interact respectively with 35 and 47 proteins involved in RNA catabolic processes (Figure 7A, yellow), RNA export (Figure 7A, light red), RNA processing (Figure 7A, blue) and/or RNA translation (Figure 7A, green), respectively. RNA processing factors interacting with Tax and HBZ are categorized into pre-mRNA processing and splicing factors (Figure 7B). To further explore the physiological roles for Tax- and HBZ-dependent effects on host mRNA splicing, we first performed motif enrichment analysis of alternative splicing events detected in patients with ATLL. We found significant enrichment for 31 RNA-binding motifs including the complementary factor for APOBEC-1 (A1CF), an RNA-binding protein regulating metabolic enzymes via alternative splicing (Nikolaou et al., 2019), identified here as a HBZ partner, and the U2 small nuclear ribonucleoprotein particle (snRNP) auxiliary factor (U2AF) large subunit U2AF65 (also called U2AF2), identified here as a Tax partner (Figures 7C, S5A and Table S1).

**Figure 7.**
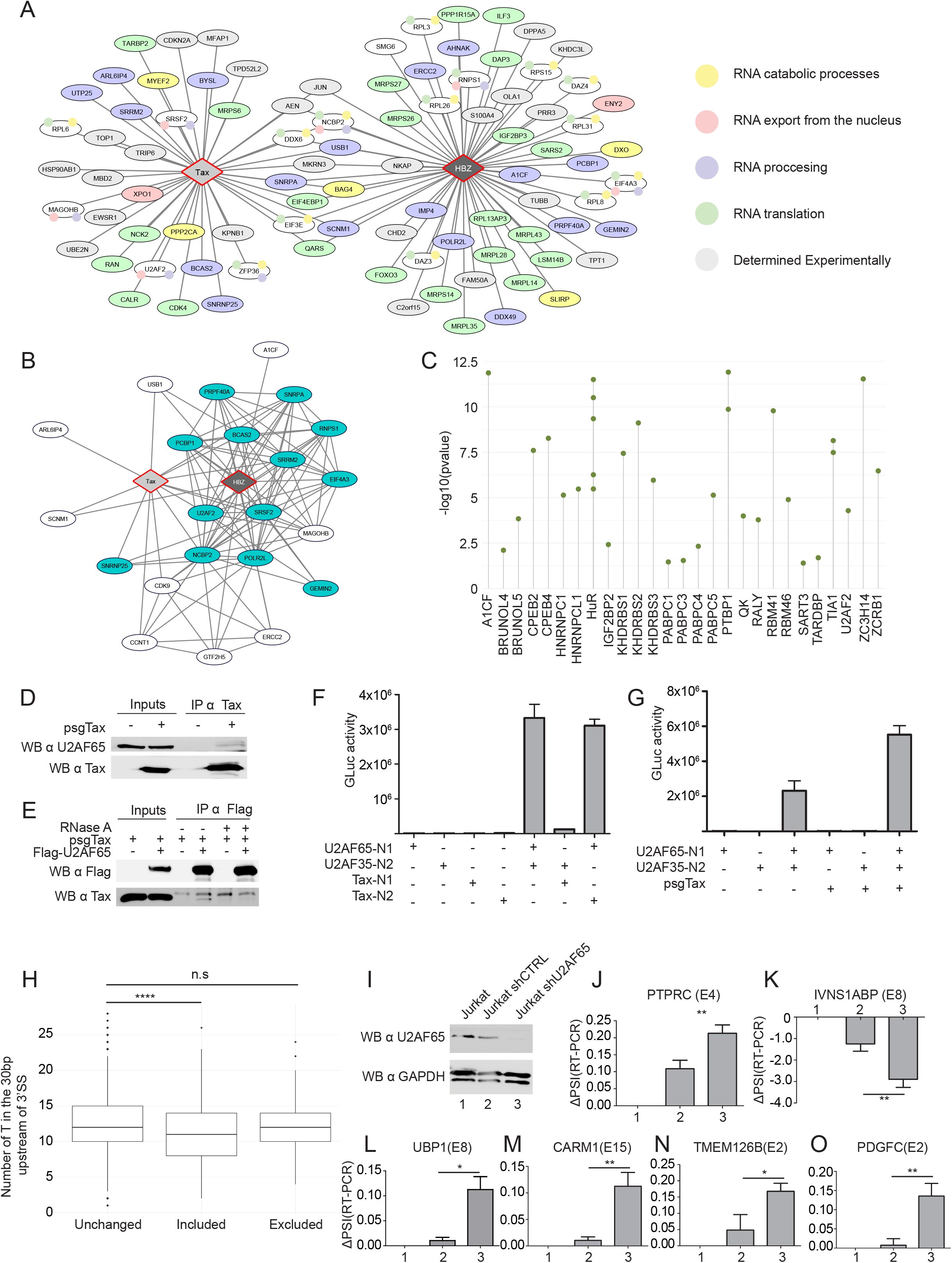
Identification of RNA splicing-specific roles for Tax and HBZ proteins. (A)Interactome of Tax and HBZ with PTRs, color-coded according to their GO categories. (B)Proteins involved in splicing (blue) are represented by wide edges. Thinner edges represent interactions between host proteins. **(C)** RNA-binding protein motif enrichment analysis of ASEs detected in ATLL samples. **(D)** Co-Immunoprecipitation of Tax and U2AF65 in HEK293T cells. **(E)** Co-Immunoprecipitations of U2AF65 and Tax, in cells treated or not with RNAse A. **(F)** GPCA assay for Tax and U2AF complex subunits. Y-axis shows average luciferase activities of three independent experiments. **(G)** GPCA assay for U2AF complex subunits under expression of Tax. Y-axis shows average luciferase activities of three independent experiments. **(H)** Number of T in the 30bp upstream of the 3’SS of alternatively spliced exons in Tax expressing cells. P = 4.044e-08. **(I)** Immunoblotting of U2AF65 in Jurkat cells knocked down for *U2AF65* expression and control cells. **(J-O)** ΔPSI of AS exons in both conditions. Regulated exons are indicated in brackets. See also Figure S5.

The U2AF complex is a well-established essential component of the spliceosome assembly pathway (Guth et al., 1999; Mackereth et al., 2011). U2AF65 forms a heterodimer with U2AF35 that recognizes the 3’ splice sites (3’SS) of introns (Mackereth et al., 2011). U2AF65 binds to the polypyrimidine tract (PY tract) of the intron and induces the recruitment of the U2 snRNP (Mackereth et al., 2011; Voith von Voithenberg et al., 2016). We performed co-immunoprecipitation assays and confirmed an interaction between Tax and endogenous U2AF65 (Figure 7D). This interaction was dependent on the presence of RNA (Figure 7E). Using the GPCA assay (Cassonnet et al., 2011), we confirmed direct Tax and U2AF65 interaction, and the absence of binding between Tax and U2AF35 (Figure 7F). However, Tax expression increased the formation of the U2AF35-U2AF65 heterodimer (Figure 7G) suggesting that the interaction interfaces of Tax/U2AF65 and U2AF35/U2AF65 may be different, and Tax does not disrupt, but rather stabilizes, the U2AF complex. As a control for specificity, we did not detect any interaction between the U2AF subunits and HBZ (Figure S5B).

Previous studies have shown that a more stable U2AF35/U2AF65 complex could favor the recognition of less conserved sub-optimal PY tract sequences containing a lower proportion of thymidines (T) (Guth et al., 1999; Pacheco et al., 2006).To determine if there is a global effect of the Tax/U2AF65/U2AF35 interaction on alternative splicing events, we inspected the 30bp upstream of all 3’SS of cassette exon events, whether altered or not upon Tax expression. We found a significant reduction of T content in the vicinity of 3’SS of more included exons, indicating that exons with less conserved PY tracts are more included upon Tax expression (Figure 7H). To further evaluate the interplay between Tax and U2AF65, we generated by shRNA a Jurkat T-cell line with reduced U2AF65 expression (Figure 7I). We analyzed 6 SE events detected in Jurkat-iTax cells, and as shown on Figures 7J to 7O, knockdown of U2AF65 affected splicing of all exons, and induced higher inclusion levels for 5 out 6 SE events.

In conclusion, although other RNA processing factors interacting with Tax (Figures 7A and 7B) are likely to influence the U2AF heterodimer and therefore splice site recognition, our results demonstrate that a large part of Tax-driven SE events are U2AF65-dependent, suggesting that, the stabilization of the U2AF complex by Tax potentially drives the initial steps of transcriptome diversification in HTLV-1 pathogenesis.

## DISCUSSION

Splicing events, producing multiple mRNA and protein isoforms participate in proteome diversity and contribute to phenotypic differences among cells (Barbosa-Morais et al., 2012; Wang et al., 2008). Splicing programs are often altered in cancer cells and systematic quantification of splicing events in tumors has led to the identification of cancer-specific transcripts that are translated into divergent protein isoforms participating in oncogenic processes (Barbosa-Morais et al., 2012). In the context of infectious diseases, it is well known that viruses exploit the host splicing machinery to compensate for their small genomes and expand the viral proteome (Chauhan et al., 2019). As a consequence, interactions with host RNA splicing factors have been reported for a number of viruses, including the Human Immunodeficiency (Stoltzfus, 2009), Influenza A (Thompson et al., 2018), Herpes Simplex (Bryant et al., 2001), Epstein-Barr Virus (Lee et al., 2012), Reovirus (Rivera-Serrano et al., 2017), Human Papillomaviruses (Graham and Faizo, 2017; Wu et al., 2017), human Adenovirus (Biasiotto and Akusjärvi, 2015), or human Parvovirus (Graham and Faizo, 2017). However, there is very limited understanding of the direct or indirect effects of viral products in regulating host RNA splicing.

Tax and HBZ, two HTLV1-encoded proteins that are major drivers of ATLL, have been known for many years to highjack the host gene expression programs. However, to date, they are exclusively considered as transcriptional regulators, acting at the level of mRNA synthesis. *In vivo*, while HBZ is constantly expressed in ATLL patient samples, Tax is mainly expressed in the initial steps of HTLV-1 infection and intermittently or not at all during the later stages. We thus performed high-throughput binary interactome mapping to identify novel interacting partners of Tax and HBZ and generated a homogenous expression model based on independent induction of Tax and HBZ expression in a Jurkat T-cell line. Although our experimental system does not take into account the *in vivo* dynamic interplay between both viral proteins, it does allow for a systematic and unbiased comparative analysis in order to identify Tax- and HBZ-dependent events contributing to dysregulation of cellular function.

First, we systematically identified, for Tax and HBZ viral proteins, shared and distinct human interacting partners implicated in gene expression regulation. Although we have not interrogated post-translational modification-dependent interactions, this map is the first to be reported for HTLV-1 and constitutes a valuable resource for in depth analysis of Tax and HBZ molecular functions. Our data suggest that both viral proteins interfere almost equally with all steps of mRNA life, including splicing processes, to reprogram the host cell transcriptome. Second, we used a Jurkat T-cell line model to identify potential Tax and HBZ splicing targets, which were validated in cells isolated from HTLV-1 carriers and ATLL samples (Kataoka et al., 2015). Interestingly, Tax- and HBZ-dependent splicing events affected respectively 33 and 63 genes that are also included in the Catalogue of Somatic Mutations in Cancer (COSMIC), as part of the cancer gene census (Sondka et al., 2018). One particular interesting example is the *PTPRC* gene encoding CD45, a critical regulator of immune cell development (Hermiston et al., 2003). The relative inclusion of exons 4 to 7 of *PTPRC* was modified in Tax and HBZ expressing cells, as well as in ATLL patients’ samples (Figure 6). These alternative splicing events on the pre-mRNA of *PTPRC* gene have been shown to lead to the expression of different CD45 isoforms (Streuli et al., 1987; Trowbridge and Thomas, 1994; Virts et al., 1998) dictating immune cell development (Hermiston et al., 2003). However, the mechanisms regulating CD45 isoform expression are not well understood. Protein kinase C (PKC) has been shown to induce *PTPRC* exon exclusion in a T-cell model (Lynch and Weiss, 2000), which correlates with previously shown activating mutations in *PKC* genes (Kataoka et al., 2015) and exclusion of exons 5, 6 and 7 in ATLL samples reported here (Figure 6). In order to propose an ATL-specific CD45 isoform as a potential diagnostic tool, future studies will be needed to address (i) mRNA and protein isoform expression at the single-cell level during disease progression, (ii) identification of specific extracellular ligands and possibly (iii) segregation of the functional interplay between Tax and HBZ in regulating *PTPRC* splicing.

Other examples from this study include well-described tumor-promoting genes such as *SRSF2, DNMT3A, ATM, BRCA1* and *PKM* (Barbosa-Morais et al., 2012), for which Tax- and HBZ-alternative splicing events are now associated with ATLL for the first time. Information about specific splicing isoforms of these genes would be beneficial to our understanding of ATLL biology, including perturbation of metabolic pathways. For instance, *PKM* pre-mRNA exhibits MXE changes in HTLV carriers (Tables S3). The encoded enzyme, pyruvate kinase PKM, is essential in glycolytic ATP production (Israelsen and Vander Heiden, 2015). Furthermore, we found that pre-mRNAs whose splicing is modified following HBZ expression are significantly enriched in metabolic processes (Tables S3). Lastly, we showed that HBZ interacts with A1CF, an RNA-binding protein that regulates metabolic enzymes via alternative splicing (Nikolaou et al., 2019), and for which we observed binding motif enrichment in ATLL samples (Figure 7C). We propose that HBZ, via interactions with RNA-processing factors, controls metabolic pathways leading to the maintenance of infected cells in a low glucose concentration and highly hypoxic host microenvironment, as recently described (Kulkarni et al., 2017). It will be interesting to explore whether inhibitors/activators of HBZ interactions with specific splicing factors targeting metabolic pathways may reveal novel therapies for ATLL.

### Concluding remarks

Just as the identification of major transcription factors interacting with Tax and HBZ (CREB/ATF, AP-1, β-catenin, Smad and NF-kB) revolutionized our understanding of the mechanisms underlying HTLV-1 pathogenesis, our finding that both viral proteins are also able to manipulate post-transcriptional events provides an excellent opportunity for an in-depth analysis of less-explored gene expression events (RNA catabolism, export, splicing and translation).

Of broad interest, our work is a contribution toward the cancer genome atlas exploration that has already revealed alternative splicing events across 32 cancer types and highlighted the U2AF complex as a key player in cancer transcriptome diversification (Rahman et al., 2013; Kahles et al., 2018). As shown here, the retroviral proteins Tax and HBZ are excellent tools to analyze transcriptional dynamics in cancer cells, beyond the evaluation of mutated genes. We advance the hypothesis that the viral protein Tax could reprogram initial steps of the T-cell proteome by hijacking the U2AF complex function. In fact, the U2AF complex targets the 3’ splice site of ∼88% of protein coding transcripts (Rahman et al., 2013), making it a perfect target for a transforming oncovirus.

## STAR METHODS

Detailed methods are provided in the online version of this paper and include the following:

### KEY RESOURCES TABLE

#### LEAD CONTACT AND MATERIALS AVAILABILITY

##### EXPERIMENTAL MODELS

° Yeast strains
° Bacterial strains
° Cell lines

### METHODS DETAILS

° Generation of Jurkat cell lines with inducible Tax or HBZ expression
° Generation of Jurkat shU2AF65
° Cell culture, transfections and treatments
° RNA extraction, RT-PCR and RT-qPCR
° Immunofluorescence and confocal microscopy
° Transmission electron microscopy
° Cloning and plasmids
° Yeast two-hybrid assay
° Immunoprecipitation and immunoblotting
° Mini gene reporter assay
° Protein complementation assay

### QUANTIFICATION AND STATISTICAL ANALYSIS

° RNA-seq data analysis
° Gene enrichment analysis
° RNA-binding motif enrichment analysis
° Statistical analysis

## DATA AVAILABILITY

## ACKNOWLEDGMENTS

We thank Dr. Clara L. Kielkopf (University of Rochester Medical Center, USA) and Dr. Yves Jacob (Institut Pasteur, Paris, France) for DNA constructs. Computational resources have been provided by the Consortium des Équipements de Calcul Intensif (CÉCI), funded by the Fonds de la Recherche Scientifique (FRS-FNRS, Belgium) under Grant No. 2.5020.11 and by the Walloon Region. This work was primarily supported by the FRS-FNRS grants PDR 14461191 and Televie 30823819 to J-C.T; Fund for Research Training in Industry and Agriculture grants 24343558 and 29315509 to C.V.; Flanders Research Foundation grant # G0D6817N and KU Leuven grant (“Vaast Leysen Leerstoel”) to J.V.W; and National Institutes of Health grants P50HG004233 to M.V. and U41HG001715 to M.V., D.E.H., and M.A.C.

## AUTHOR CONTRIBUTIONS

Conceptualization, C.V., J-C.T., and F.D.; Methodology, C.V., T.OG., A.D., B.C., F.M., L. B.A. and J-C.T; Investigation, C.V., B.G., M.C., M.T. and J-C.T.; Formal analysis, C.V., T.OG., B.C., G.C., L.B.A., Y.V.W., M.T. and F.M.; Resources, K.K., S.O., M.A.C, D.E.H., and M.V.; Writing - Original Draft, C.V. and J-C.T.; Writing - Review and Editing, J-C.T., F.D., T.OG., J.O., J.V.W, F.M., M.A.C, D.E.H., Y.J. and M.V; Supervision, J-C.T., M.V. and F.D.; Funding Acquisition, C.V., J-C.T., F.D., Y.V.W., D.E.H.,M.A.C. and M.V.

## DECLARATION OF INTERESTS

The authors declare no competing interests

## STAR METHODS

### KEY RESOURCES TABLE

| REAGENT or RESOURCE | SOURCE | IDENTIFIER |
| --- | --- | --- |
| **Antibodies** |  |  |
| Anti-SC35 | Abcam | PRID: AB_298608 |
| Anti-Flag M2 | Sigma-Aldrich | PRID: AB_259529 |
| Anti-Flag (Rabbit) | Sigma-Aldrich | PRID: AB_439687 |
| Anti-Tax | Francoise Bex | Meertens et al., 2004 |
| Anti-GAPDH | Santa Cruz | PRID: AB_2107299 |
| Anti-HA | Santa Cruz | PRID: AB_627809 |
| Anti-myc | Santa Cruz | PRID: AB_2857941 |
| **Chemicals** |  |  |
| Doxycycline hydrochloride | Fisher Scientific | Cat#10592-13-9 |
| Protamine sulfate | MP Biomedicals | Cat#9009-65-8 |
| Blasticidin S | Sigma-Aldrich | Cat# 15205 |
| Hygromycin B | Sigma-Aldrich | Cat# 10843555001 |
| **Deposited Data** |  |  |
| RNA-seq Raw and processed data | This paper | GEO: GSE146210 |
| **Experimental Models: Cell Lines** |  |  |
| HeLa | ATCC | CRM-CRL-2 |
| HEK293T | ATCC | CRL-3216 |
| Hek293 iHBZ | This paper | N/A |
| Jurkat | ATCC | CSC-C9455L |
| Lenti-X-293T cell | Clontech-Takara | Takara #632180 |
| Jurkat Tet3G iTax and Jurkat Tet3G iHBZ | This paper | N/A |
| Jurkat-iTax and Jurkat-iHBZ | This paper | N/A |
| **Experimental Models: Organisms/Strains** |  |  |
| *S.cerevisiae*: Strain background: MATa Y8800 | (Yu et al., 2008) | N/A |
| *S.cerevisiae*: Strain background: MATa Y8930 | (Yu et al., 2008) | N/A |
| Recombinant DNA |  |  |
| pLenti-CMVtight-Blast-DEST | Eric Campeau | Addgene#26434 |
| pLenti CMV rtTA3 | Eric Campeau | Addgene #26429 |
| pLVX-Tet3G | Clontech-Takara | Takara Cat#631187 |
| psPAX2 | Didier Trono | Addgene # 12260 |
| pCMV-VSVG | (Stewart, 2003) | Addgene #8454 |
| pLenti6 Tight-Tax | This paper | N/A |
| pLenti6 Tight-Flag-HBZ | This paper | N/A |
| pLV U6 shRNA hU2AF2 (mCherry-Neo) | This paper | N/A |
| humanORFeome v7.1 and v8.1 | The Center for Cancer Systems Biology (CCSB-DFCI) | https://ccsb.dana-farber.org/resources.htm |
| pDEST-AD-CEN SY | (Dreze et al., 2010) | N/A |
| pDEST-DB-CEN SY | (Dreze et al., 2010) | N/A |
| pGLucN1 and pGLucN2 | (Cassonnet et al., 2011) | N/A |
| Software and Algorithms |  |  |
| ImageJ | (Schneider et al., 2012) | https://imagej.nih.gov/ij/ |
| STAR | (Dobin et al., 2013) | https://code.google.com/archive/p/rna-star/ |
| DESeq2 | (Love et al., 2014) | http://bioconductor.org/packages/release/bioc/html/DESeq2.html |
| rMATS | (Shen et al., 2014) | http://rnaseq-mats.sourceforge.net/ |
| GSEA 3.0 | (Liberzon et al., 2015) | https://software.broadinstitute.org/gsea/index.jsp |
| GORILLA | (Eden et al., 2009) | http://cbl-gorilla.cs.technion.ac.il/ |
| AME | (McLeay and Bailey, 2010) | http://meme-suite.org/ |

### CONTACT FOR REAGENT AND RESOURCE SHARING

CONTACT FOR REAGENT AND RESOURCE reagents should be directed to and will be fulfilled by the Lead Contact, Dr. Jean-Claude Twizere.

Plasmids and cell lines generated in this study are available upon request and approval of the Material Transfer Agreement (MTA) by the University of Liege.

### EXPERIMENTAL MODEL AND SUBJECT DETAILS

### EXPERIMENTAL MODEL AND SUBJECT DETAILS

#### Yeast strains

Yeast strains used in this study are derived from S288C as described (Dreze, et al 2010). Haploids of both mating type MATα Y8930C and MATa Y8800C were used. These strains harbor the following genotype: leu2-3,112 trp1-901 his3Δ200 ura3-52 gal4Δ gal80Δ GAL2::ADE2 GAL1::HIS3@LYS2 GAL7::lacZ@MET2cyh2R. Yeast cells were cultured either in selective synthetic complete (SC) media or in rich medium (YPD) supplemented with glucose and adenine. Cells were incubated at 30°C.

#### Bacterial strains

Chemically competent DH5α E. coli cells were used for all bacterial transformation in this study. Post transformation, cells were cultured in LB (25 g/l) supplemented with antibiotics (50 µg/ml of ampicillin, spectinomycin or kanamycin) and incubated at 37°C for 24 hours.

#### Cell lines

HEK293T (*Homo sapiens*, fetal kidney) and HeLa (*Homo sapiens*, Henrietta Laks cervical cancer) cells were cultured in Dulbecco’s Modified Eagle Medium (DMEM) supplemented with 10% fetal bovine serum, 2mmol/L L-glutamine and 100 I.U./mL penicillin and 100μg/mL streptomycin. Cells were incubated at 37°C with 5% CO2 and 95% humidity. Jurkat (*Homo sapiens*, T-cell leukemia) cells were cultured in Roswell Park Memorial Institute (RPMI) media supplemented with 10% fetal bovine serum, 2mmol/L L-glutamine and 100 I.U./mL penicillin and 100 μg/mL streptomycin. Cells were incubated at 37°C with 5% CO2 and 95% humidity.

### METHOD DETAILS

#### Yeast two hybrid assay

Details of the Y2H screening method are described elsewhere (Dreze et al., 2010; Bergiers et al., 2014). Y2H assays were performed with 19 distinct fragments of the Tax protein and 9 fragments of HBZ protein cloned in CEN plasmids pDEST-AD-*CYH2* (AD vector) and pDEST-DB (DB vector). AD and DB vectors were transformed into MATa Y8800 and MAT*α* Y8930 *Saccharomyces cerevisiae* strains, respectively (Dreze et al., 2010; Yu et al., 2008). A total of 3652 Human ORFs, corresponding to human transcription factors and RNA-binding proteins, were obtained from the human ORFeome v7.1 collection. For Y2H screening, we pooled MATa Y8800 yeast strains containing human and viral AD-ORFs, and performed mating against MAT*α* Y8930 yeast clones containing individual human or viral DB-ORFs. After selection of mated yeasts on medium lacking leucine, tryptophan and histidine, the identity of the interacting protein pairs was determined by sequencing of the corresponding ORF clone. A counter-selection on medium containing cycloheximide was performed simultaneously in the screens to identify false positives (Walhout & Vidal, 1999: Dreze, et al). All discovered interacting pairs were retested again similarly but in pairwise screenings. Interacting pairs confirmed in the second screening were considered positives. Construction of interacting networks was done with Cytoscape v3.5.1.(Shannon, 2003).

#### Generation of Jurkat cell lines with inducible Tax or HBZ expression

The viral Tax gene and the FLAG-tagged HBZ gene were cloned into the pLenti-CMVtight-Blast-DEST (w762-1) vector from Addgene (#26434) with the Gateway cloning system. Production of lentiviral vectors (2nd generation) and generation of the Jurkat Tet3G iTax and Jurkat Tet3G iHBZ cell lines was performed by the GIGA Viral Vectors Platform at the University of Liège with the 3^*rd*^generation inducible gene expression system Tet-On (from Clontech-Takara).The plasmids pLVX-Tet3G, pLenti6 Tight Tax and pLenti6 Tight Flag-HBZ were transfected separately on *Lenti*− *X™*293T Cell Line (from Clontech-Takara) each with a lentiviral packaging mix. This mix contained a packaging construct with the gag, pol, rev genes (psPAX2), and an Env plasmid expressing the vesicular stomatitis virus envelope glycoprotein G (VSV-G). The lentiviral supernatants were harvested, concentrated and tittered by qRT-PCR (Lentiviral Titration Kit LV900 from AbmGood) to be further used for transduction. The Jurkat cell lines (5 × 10^5^ cells/ml) were co-transduced with lentiviral vectors pLVX-Tet3G at a multiplicity of infection (MOI) of 30 and with pLenti6 Tight-Tax or -Flag-HBZ at a MOI of 16. For this transduction step, the reagent protamine sulfate (MP Biomedicals) was used according to the manufacturer’s instructions (8 *µ*g/ml). Subsequently, the cells were centrifuged at 800 x g for 30 minutes at 37 degrees. The pellet was suspended in RPMI-1640 containing 10% FBS. After 72h cells were cultivated in cell culture medium containing blasticidin(10 *µ*g/ml) and hygromycin B (400 *µ*g/ml) in order to select transduced cells expressing the gene of interest (Tax or Flag-HBZ) and the rtTA3 gene until the non-transduced cells (negative control) died as determined by Trypan Blue staining.

#### Generation of Jurkat shU2AF65 cells

Hek293 Lenti-x 1B4 cells (Clontech) were transfected with pLV U6 shRNA hU2AF2 (mCherry-Neo) (VectorBuilder), pVSV-G (Clontech, PT3343-5) and psPAX2 (Addgene, 12260). Supernatants were harvested and viruses were concentrated by ultracentrifugation. Jurkat cells were transduced with viruses at a MOI of 30. After 72h cells were selected with 2 mg/ml neomycin (G418, Invitrogen).

#### Cell culture, transfections and treatments

Induction of Tax or Flag-HBZ in Jurkat-iTax or -iHBZ cells was performed by treating cells with doxycycline hydrochloride (1 µg/ml, Fisher Scientific) for 48h. HEK293T and HeLa cells were transfected using polyethylenimine (PEI25K, Polysciences) (from 1 mg/ml) by a ratio 2:1 to plasmid concentration. HEK293T and HeLa cell lines were maintained in DMEM (Gibco) complemented with 10% FBS while Jurkat cell lines were maintained in RPMI (Gibco) also complemented with 10% FBS. Cell lines were regularly checked for mycoplasma contamination.

#### RNA extraction, RT-PCR and RT-qPCR

Total RNA extraction from cell pellets was performed according to the manufacturer’s protocol (Nucleo Spin RNA kit from Macherey-Nagel) and cDNA was obtained by reverse transcription with random primers using the RevertAid RT Reverse Transcription Kit from Thermo Fisher. One or half a microgram of total RNA was used to make cDNA, which were diluted 100 times to perform PCR amplification with specific primers, using Taq polymerase (Thermo Fisher). PCR products were migrated by SDS-PAGE and revealed with Gel Star (Lonza) under UV light. Bands were quantified using ImageJ software. Quantitative PCR (qPCR) reactions were done with iTaq Universal SYBR Green Supermix (BioRad) in triplicates on a Light Cycler 480 instrument (Roche). The ΔΔCt method was used to analyze relative target mRNA levels with GAPDH as an internal control. All primers used in the study are presented in Table S4.

#### Immunofluorescence and confocal microscopy

Transfected HeLa cells with GFP-HBZ or an empty plasmid were grown on glass coverslips for 24h. Cells were washed in PBS and fixed with 4% paraformaldehyde (PAF) for 15 min. After washing with PBS, cells were permeabilized in PBS with 0.1% Triton X-100 for 5 min. Cells were then incubated in blocking solution (PBS with 4% BSA)for 1h before incubation with anti-SC35 primary antibody (Abcam) overnight at 4 degrees. Samples were then incubated with Alexa568-conjugated secondary antibodies (Thermo Fisher) and further incubated 10 min. with DAPI (Thermo Scientific, in PBS) before washing and mounting with Mowiol. All images were acquired with a Nikon A1 confocal microscope and processed with ImageJ.

#### Transmission electron microscopy

Hek293 cells inducible for HBZ expression (Hek293 iHBZ) were generated similarly to Jurkat-iTax and -iHBZ cell lines. However a plasmid pLenti CMV rtTA3 was used instead of the plasmids pLVX-Tet3G. HBZ expression was induced or not in Hek293 iHBZ cells for 48 hours with doxycycline before cells were fixed for 90 minutes at 4 degree C with 2.5% glutaraldehyde in a Sörensen 0.1 M phosphate buffer (pH 7.4) and post-fixed for 30 min with 2% osmium tetroxide. After dehydration in graded ethanol, samples were embedded in Epon. Ultrathin sections obtained with a Reichert Ultracut S ultra microtome were contrasted with uranyl acetate and lead citrate. Observations were made with a Jeol JEM-1400 transmission electron microscope at 80kV.

#### Cloning and plasmids

Viral clones were amplified by PCR using specific primers flanked at the 5’ end with AttB1.1 and AttB2.1 Gateway sequences and were inserted into pDONR223 by Gateway cloning (Invitrogen). Human ORFs (encoding U2AF65, SNRPA, eIF4A3) were retrieved from the human ORFeome v7.1 or the human ORFeome v8.1 (Yang, X, 2011). Inserts in pDONR223 were then transferred into appropriate destination vectors by LR cloning. A list of Tax and HBZ clones is available in Table S4.

#### Cell lysates, immunoprecipitation, immunoblotting and antibodies

For Immunoprecipitations (IPs) HEK293T cells were harvested and lysed in IPLS (Immuno Precipitation Low Salt; Tris-HCl pH 7.5 50 mM, EDTA, pH 8, 0.5 mM, 0.5% NP-40, 10% glycerol, 120 mM NaCl, cOmplete Protease Inhibitor (Roche)). RNaseA treatment (Thermo Fisher Scientific, 10 *µ*g/ml at 37% for 30 min) was performed on cleared lysates when indicated. Supernatants were incubated with anti-FLAG M2 agarose beads (Sigma-Aldrich) and then were washed with IPLS (incubation times and number of washes depended on tested interactions). For semi-endogenous IPs, rabbit anti-Tax antibody (Meertens et al., 2004) was incubated overnight with cell lysates. Afterwards, Protein A/G PLUS-Agarose beads (Santa Cruz) were incubated with lysates for 2h and beads were washed 3 times with IPLS. Beads were re-suspended in 2x SDS loading buffer and boiled. Samples were then analyzed by SDS-PAGE and western blotting and revealed with ECL detection kit (GE Healthcare Bio-Sciences) according to standard procedures.

#### Protein Complementation Assay (PCA)

Tax, HBZ and interacting partners ORFs were cloned in destination vectors containing GLucN1 and GLucN2 fragments of the *Gaussia princeps* luciferase. HEK293T cells were seeded in 24-well or 96-well plates and transfected with 500 ng or 200 ng of the appropriate constructs (GLucN1 + GLucN2), respectively. After 24h cells were washed with PBS and lysed using the manufacturer’s lysing buffer (Renilla luciferase kit, Promega). Ten to 20 *µ*l of lysates were then used to quantify luminescence in a Centro lb 960 luminometer (Berthold).

### QUANTIFICATION AND STATISTICAL ANALYSIS

#### RNA-seq data analysis

Libraries were prepared with the Illumina Truseq stranded mRNA sample prep kit and paired-end sequencing was performed with the Illumina NextSeq500 PE2X75 system by the Genomics platform at GIGA, University of Liege. Sequence reads were aligned to the human genome hg19 (UCSC) using STAR (Dobin et al., 2013). Differential expression analysis was performed with DESeq2 (Love et al., 2014) on read counts from STAR quant Mode. Genes were considered significantly up- or down-regulated if their base 2 logarithm fold change was *>*1 or *<*-1 and their adjusted p-value was *<*0.01. Analysis of Alternative Splicing Events (ASEs) was performed with the rMATS software v3.2.1 (Shen et al., 2014). Genes with low expression levels were removed from the results by filtering out genes with a TPM *<*1. TPM was calculated using Salmon v0.9.1. ASEs were further filtered to consider only events with a ΔPSI *>*1 or *<*-1 and with a FDR *<*0.05. Sashimi plots were generated using rmats2sashimi. RNA-seq data from patients were previously described (Kataoka et al., 2015).

#### Gene enrichment analysis

Gene enrichment analysis on differentially expressed genes was performed with GSEA 3.0 (Gene Set Enrichment Analysis) in pre-ranked mode using the hallmark gene sets from the Molecular Signatures Database (Liberzon et al., 2015). GO enrichment analysis was performed on alternatively spliced genes with Gorilla(Eden et al., 2009)using all detected ASEs as a background. Other enrichment analyses were performed by hyper geometric tests or by calculating empirical p-values in R.

### RNA-Binding motif enrichment analysis

Sequences of regulated exons detected by rMATS (SE events only) and up to 200 bp of their flanking introns were retrieved using bed tools v0.9.1. Control sequences consisted of detected ASEs with low ΔPSI values and high FDR. The FASTA files generated were then interrogated for known RNA-binding motifs from the literature (Ray et al., 2013) using the AME software from MEME suite 5.0.2 with default settings (McLeay and Bailey, 2010). Motifs with an adjusted p-value *<*0.05 were considered significant.

## DATA AVAILABILITY

RNA-sequencing data have been deposited in NCBI’s Gene Expression Omnibus (Edgar, 2002) and are accessible through GEO accession number GSE146210 (https://www.ncbi.nlm.nih.gov/geo/query/acc.cgi?acc=GSE146210).

---

**Figure S1.**
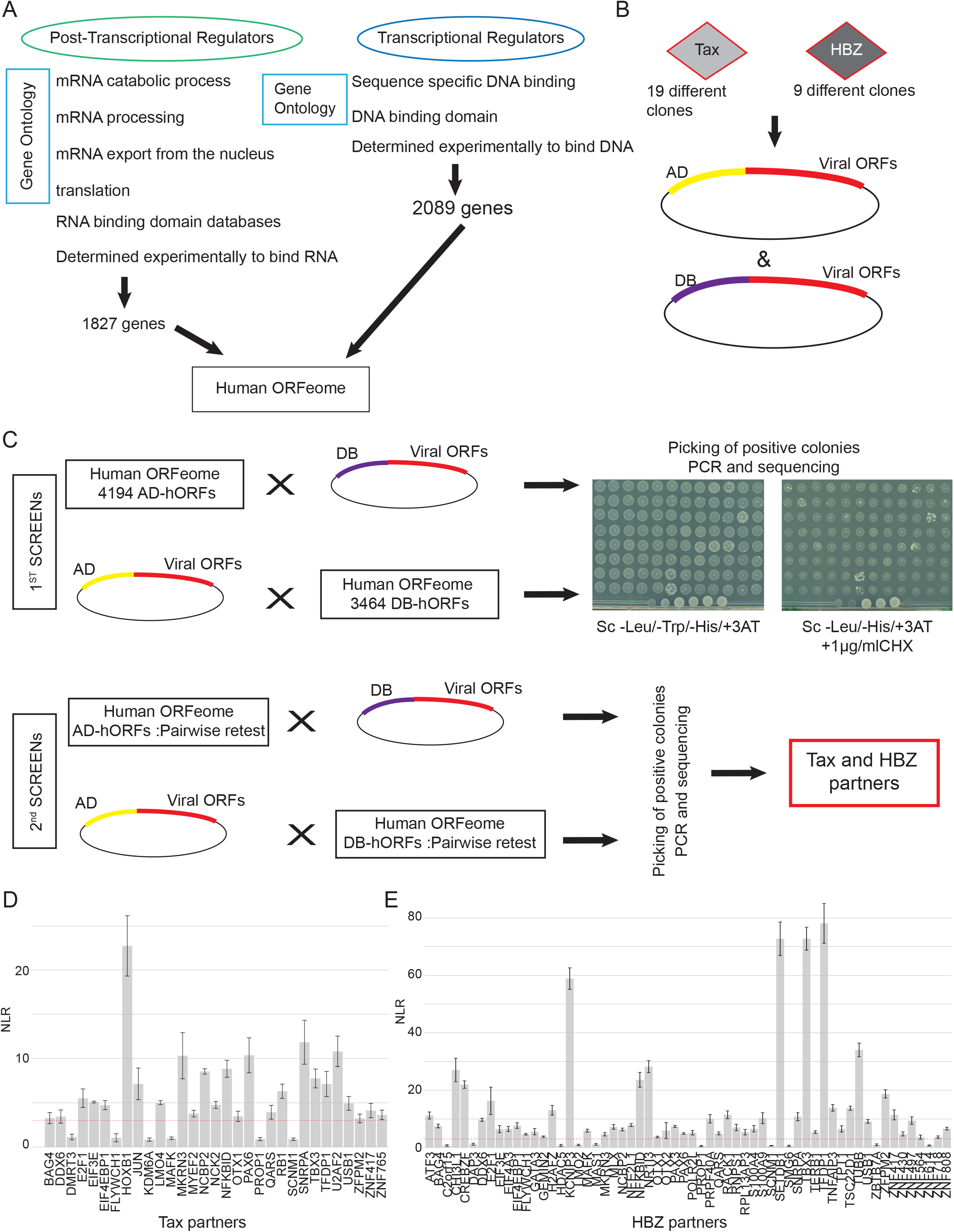
Overview of the Yeast-two hybrid (Y2H) experiment. Related to Figure 1. **(A)**. Pipeline to generate a comprehensive list of human RBPs and TFs. **(B)** Cloning strategy for the Tax/HBZ mini-library. **(C)** Y2H strategy to identify high-quality protein-protein interactions (PPIs).

**Figure S2.**
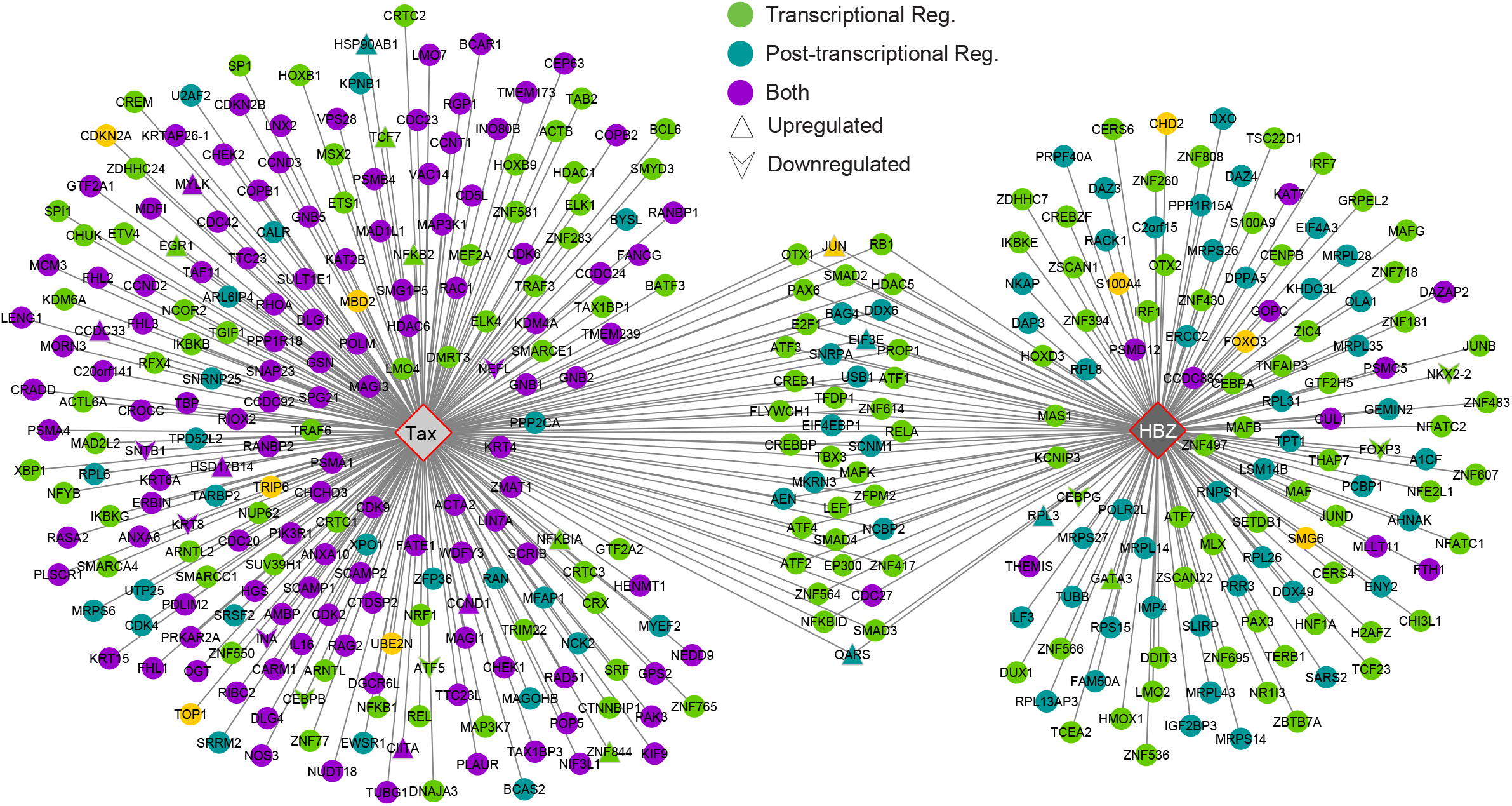
A comprehensive map of host proteins interacting with Tax and HBZ. Related to Figure 1. Host proteins are color-coded according to their function in Transcriptional Regulation (TR), Post-Transcriptional Regulation (PTR) or other (purple). Upward triangles and downward arrows show genes with an up-regulation or down-regulation following Tax or HBZ expression, respectively.

**Figure S3.**
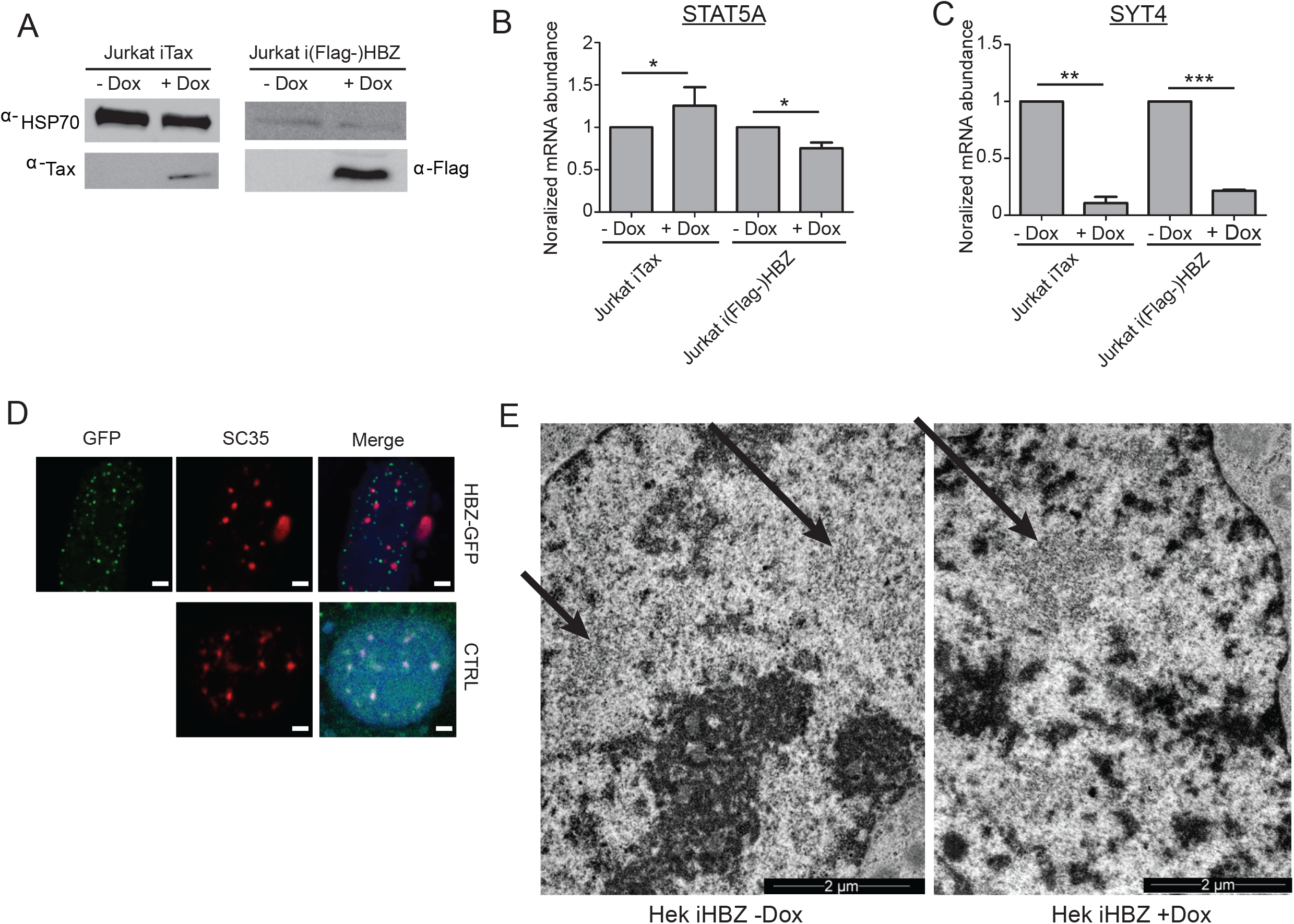
Perturbation of nuclear speckles and interchromatin granules by Tax and HBZ. Related to Figure 3. **(A)** Western blot showing expression of Tax or HBZ in Jurkat-iTax or -iHBZ cells, respectively, upon induction by doxycycline for 24 hours at 1mg/ml. **(B-C)** qRT-PCR analysis showing variation of normalized mRNA abundance of *STAT5* A **(B)** and *SYT4* **(C)** upon expression of Tax and HBZ. **(D)** Immunofluorescence microscopy indicates that SC35, a marker of nuclear speckles, localizes into more round shapes upon expression of HBZ in HEK293T cells. Scale bars = 2 *µ*m. **(E)** Partial nucleus of HEK293T iHBZ cells induced for HBZ expression (left) or not (right) by TEM. Interchromatin granules (also known as nuclear speckles) are indicated by black arrows and display a more compact phenotype upon HBZ expression. Scale bars = 2 *µ*m.

**Figure S4.**
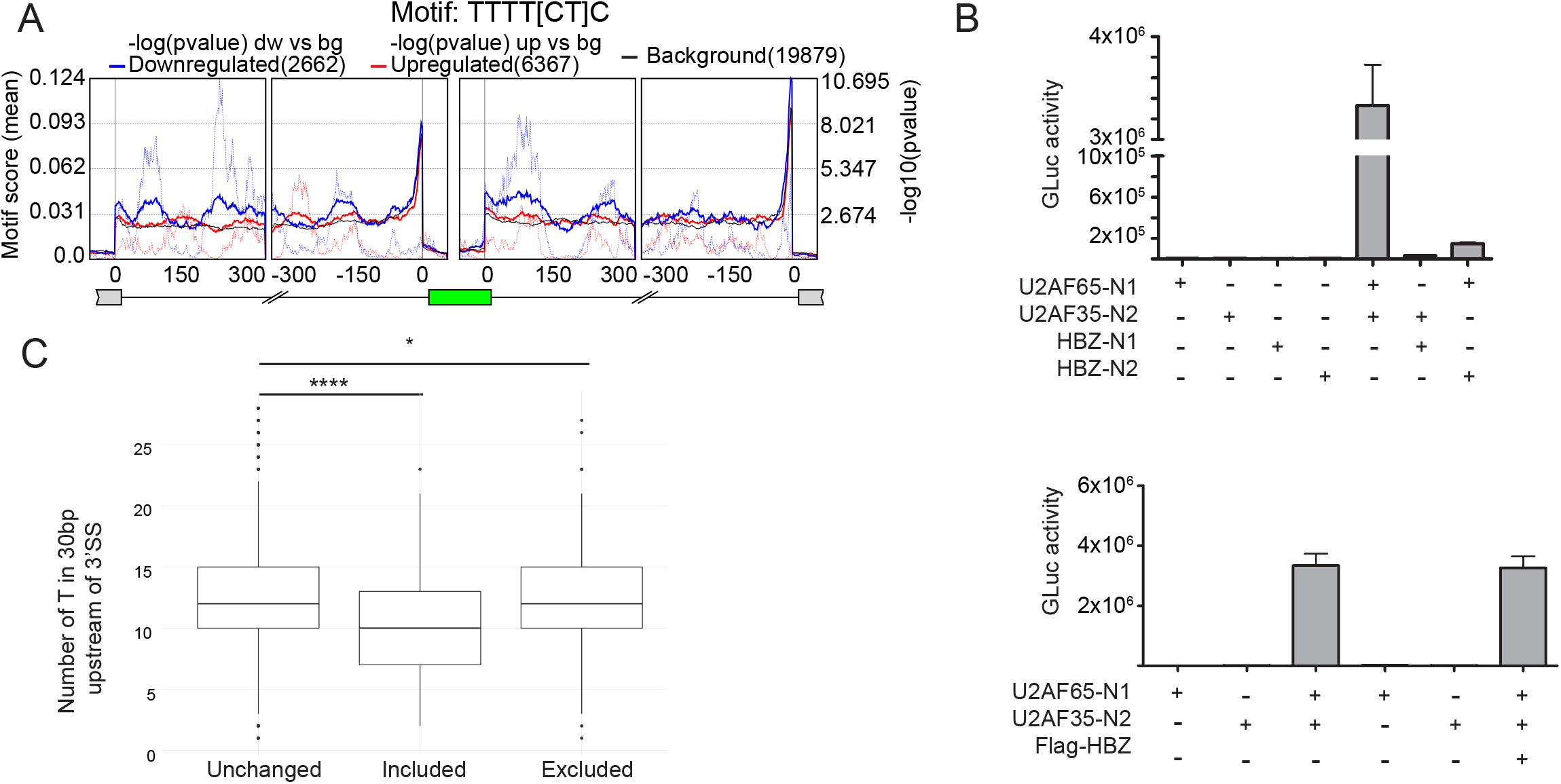
Alternative splicing in ATLL patient samples. Related to Figure 4. Table of alternatively spliced genes and their affected exons in ATLL patients, Jurkat-iTax and Jurkat-iHBZ cells

**Figure S5. Spatial distribution of U2AF2 binding-motif in ATLL patients. Related to Figure 5**

**(A)** Solid lines indicate the mean U2AF2 binding motif score calculated in a 50 bp sliding window. Dotted lines indicate -log10 p-values obtained by statistical comparison of motif scores between modified exons (exclusion=down-regulated and inclusion=up-regulated) against non-modified background exons. Green box represents regulated exons flanked by neighboring introns and upstream and downstream exons in black lines and grey boxes. **(B)** GPCA to test the absence of interaction between HBZ and U2AF complex subunits (U2AF65 and U2AF35). Y-axis shows luciferase activity for a representative experiment of 3 repetitions. **(C)** Number of Ts in the 30bp upstream of 3’SS of alternatively spliced exons (SE events) in Jurkat-iHBZ cells. Included P *<*2.2e-16, Excluded P = 0.03398, by Welch Two Sample t-test in R.

Table of alternatively spliced genes and their affected exons in ATLL patients, Jurkat iTax and iHBZ cells

| Gene name | Alternatively spliced Exon | Main known functions | Feature on exon | References |
| --- | --- | --- | --- | --- |
| **MSMO1** | SE event 2 | Oxidoreductase | Ubiquitinated lysine<br>Nuclear localization signal<br>Helical transmembrane region | FasterDB<sup>1</sup>, pfam |
| **ILF2** | SE event 12 | T-cell activation<br>DNA binding | DZF domain (unknown function) | FasterDB<sup>1</sup>, pfam |
| **PTPRC (CD45)** | SE events 4,5,6,7 | T cell activation | Intrinsically unstructured polypeptide region Cysteine rich domain<br>O-glycosylation site<br>Fibronectin like domains | FasterDB<sup>1</sup>, pfam<br>Tchilian and Beverly, 2006<sup>2</sup> |
| **ATF2** | SE event 14 | Transcription factor | Premature Stop codon | FasterDB<sup>1</sup>, pfam |
| **EYA3** | SE event 13 | Tyrosine phosphatase<br>DNA repair | Premature Stop codon | FasterDB<sup>1</sup>, pfam |
| **IVNS1ABP** | SE event 8 | RNA splicing<br>Viral processes | BTB/Kelch-associated domain | FasterDB<sup>1</sup>, pfam |
| **VPS13C** | SE event | Mitochondrial function |  | FasterDB<sup>1</sup>, pfam |
| **PCNP** | A3'SS event 2 | Protein binding<br>Cell cycle | PTMs<br>Intrinsically unstructured polypeptide region | FasterDB<sup>1</sup>, pfam |
| **WBSCR22** | A5'SS 2 | rRNA methyltransferase activity | Intrinsically unstructured polypeptide region | FasterDB<sup>1</sup>, pfam |
| **WTAP** | RI 7-8 | mRNA splicing<br>Cell cycle | Ubiquitinated lysine | FasterDB<sup>1</sup>, pfam |

<sup>1</sup> http://fasterdb.ens-lyon.fr/
<sup>2</sup> Tchilian EZ, Beverley PC, "Altered CD45 expression and disease", *Trends Immunol.*, vol. 27, pp. 146-153, Mar 2006.

## Notes

### Competing Interest Statement

The authors have declared no competing interest.

